# Pathway Thermodynamic Analysis Postulates Change in Glutamate Metabolism as a Key Factor in Modulating Immune Responses

**DOI:** 10.1101/2024.12.06.627255

**Authors:** Sunayana Malla, Rajib Saha

## Abstract

**Background:** Temperature, as seen during fever, plays a pivotal role in modulating immune responses and maintaining cellular homeostasis. Shifts in temperature influence the thermodynamic feasibility of metabolic reactions, with Gibbs free energy (ΔG) serving as a key indicator of the spontaneity of reactions under specific conditions. By altering ΔG in response to temperature changes across various metabolite concentrations and cell types, we can gain insights into the thermodynamic properties of metabolic pathways and identify critical factors involved in metabolism and immune function. Using Max-Min Driving Force (MDF) analysis, we can assess changes in ΔG by varying temperature and metabolite concentrations, allowing for a detailed examination of thermodynamic feasibility at both the pathway and individual reaction levels.

**Results:** In this study, MDF analysis is applied to measure the changes in the driving force of pathways and the ΔG of each reaction at normal human core temperature (310.15 K) and elevated temperatures (up to 315.15 K). Additionally, we explore how shifts in the thermodynamic feasibility of reactions under immune activation, compared to normal physiological conditions, highlight key metabolic intermediates—such as fructose-1,6-bisphosphate, glucose-6-phosphate, and several steps in glutamate metabolism—as important regulators of metabolic processes and immune responses.

**Conclusion:** The goal of this study is to underscore the value of thermodynamic parameters such as ΔG, concentration, and temperature in identifying potential therapeutic targets, with the aim of mitigating the detrimental effects of fever while preserving its beneficial aspects.

## Background

Temperature fluctuations, such as fever (hyperthermia), have profound effects on the immune system^1^ and cellular homeostasis. The immune system is susceptible to environmental changes, and temperature shifts can directly affect the metabolism, activation, and efficiency of cells, such as macrophages, neutrophils, erythrocytes, and dendritic cells^2^. At the molecular level, temperature alters the physical properties of cellular membranes, protein stability, and enzyme activity which cascades into broader effects on immune cell metabolism, such as folding of proteins and expression of heat shock proteins^3^. For a comprehensive understanding of these phenomena, thermodynamic analysis provides essential insights into the underlying physical chemistry that governs metabolic and immune processes. Reactions in immune cells must have sufficient driving forces, represented by the Gibbs free energy (Δ*_f_*G°/ΔG°), to proceed efficiently^4^. When temperature changes, the Gibbs free energy shifts, potentially making some reactions more or less feasible. Thermodynamic models help elucidate how enzyme activity changes with temperature, affecting reaction rates within metabolic pathways^5^. Several studies have explored the thermodynamic feasibility in immune cells focusing on energetic demands, ion gradients across cell membranes, and change in ΔG° for antibodies that induce immune responses^3,6^. However, pathway thermodynamics to decipher cell capabilities at different conditions such as metabolite concentration changes and temperature fluctuations has not been fully explored yet.

Additionally, majority of the efforts to use thermodynamic parameters to study pathway and reaction feasibility are limited to simpler model organisms such as yeast and *E coli* and rarely expand to human cells^5,7^. Thermodynamic-based Metabolic Flux Analysis (TMFA) integrates thermodynamic constraints into metabolic flux models to ensure that all reactions in a metabolic network are thermodynamically feasible^8^. However, TMFA requires strict assumptions about metabolite concentrations and environmental conditions, which may not accurately reflect the actual dynamic conditions within cells, particularly in complex organisms^8,9^. These fixed assumptions can make it challenging to capture the real flexibility and adaptability of the cellular metabolism, where metabolite concentrations and environmental factors often fluctuate.^8,9^.

On the other hand, the Max/Min Driving Force (MDF) concept can identify which reactions become thermodynamically constrained or more favorable as temperatures fluctuate or physiological states change. This offers critical insights into metabolic robustness, efficiency, and adaptability, particularly for the immune cells such as neutrophils, macrophages, dendritic cells, and erythrocytes, which are often at the forefront of immune responses including oxygen transport^1^. By comparing these cell types, MDF can elucidate their metabolic adaptations, such as the high glycolytic flux in neutrophils during inflammation^10^, more active lipid metabolism in macrophages^11^, energy demands of dendritic cells during antigen presentation^12^, and erythrocytes’ reliance on anaerobic metabolism^13^. These insights provide valuable information on cell-specific vulnerabilities and pathways, informing therapeutic strategies for immune modulation and metabolic interventions in disease contexts. By focusing on maximizing the smallest driving force in a pathway, MDF can highlight reactions that become bottlenecks as temperature changes^7^. This approach is particularly critical because it not only ensures overall pathway feasibility but also pinpoints specific steps that may need adjustment or optimization in response to temperature shifts and concentration changes.

To conduct MDF analysis, accurate estimates of Δ*_f_*G° for all reactions in the pathway of interest are required^7^. Among the various databases and tools available, such as NIST^14^, MetaCyc, KEGG, and THERMDB, Equilibrator stands out due to its unique combination of features, accessibility, and versatility^15–18^. Equilibrator leverages a curated and validated thermodynamic database derived from multiple sources ensuring high reliability for biochemical reactions^7^. It also implements the Component Contribution Method (CCM), which combines experimental data with predictive models, enabling accurate Δ*_f_*G° estimates even for reactions with sparse data^4^. Additionally, Equilibrator is specifically designed for biochemical systems, unlike other tools that often focus on general chemistry or reaction mechanisms^19^. However, despite its extensive database, Equilibrator’s reliance on available experimental data limits its application only to specific pathways.^20^

Hence, we applied the MDF analysis on pathways for which we could obtain Δ*_f_*G° values from equilibrator^20^(namely glycolysis/gluconeogenesis, OXPHOS, PPP, TCA cycle, Arginine/Proline, amino sugar, and nucleotide sugar metabolism, leukotriene metabolism and other amino acid pathways), many of which were also identified by Ippolito et al, 2014 as the pathways that showed significant deviation to heat stress. We used metabolite concentrations expected in normal physiological conditions (0.01 mM to 10 mM) and further explored the pathway activity for highly essential immune cells, namely macrophages, neutrophils, erythrocytes, and dendritic cells with an increase in temperature by constraining the metabolite concentrations as reported by Hooftman et al., 2023, and Kaiser et al., 2020. We identify the reactions that show a dynamic shift in thermodynamic feasibility from each of the pathways mentioned above when the temperature increases from normal body temperature (310.15 K) to fever conditions (312.15 K) to 313.15 K −315.15 K that can be dangerous and warrant immediate medical attention^1,24,25^. The MDF analysis highlights the reactions associated with the utilization of glutamate to produce several factors such as ornithine, citrulline, and G5SSH (L-glutamate 5-semialdehyde) as significant contributors in metabolic regulation and immune responses. Overall, the results of this study add to the importance of pathway-level analysis of temperature and the benefits of using thermodynamics to identify potential biomarkers and therapeutic target-level analysis.

## Results

### Change in ΔG° with Temperature Increase

Max/Min Driving Force (MDF) analysis is a powerful tool for identifying the bottleneck and critical reactions in metabolic pathways, especially under conditions where temperature fluctuations may impact thermodynamic feasibility^7,26^. While human metabolism can adjust to minor temperature changes through homeostatic mechanisms, significant or prolonged deviations highly disrupt metabolic efficiency, enzyme function, and energy balance, ultimately compromising health^25^. Hence, examining changes in ΔG° offers unique insights into the effects of temperature fluctuations on metabolic feasibility. Since ΔG° is temperature-dependent, it directly influences the thermodynamic feasibility of metabolic reactions^27^.

To this end, we applied MDF analysis to obtain the overall driving force of pathways and ΔG° of each reaction with the substrate concentration range set between 0.01 to 10 mM. Due to the lack of experimental measurements, the 0.01 mM to 10 mM concentration range is biologically justified as it represents the metabolite concentrations levels across diverse human states, from nutrient-rich to starved conditions^7,28^. This range is widely accepted in metabolic modeling and experimental studies, allowing flexibility to capture low- and high-concentration metabolites relevant to various pathways^5,7,9^. Additionally, it aligns with established standards in systems biology for representing physiological concentrations and thus allows for a realistic assessment of how human cells adapt to temperature fluctuations^7^. We also increased the temperature from 310.15 K to 315.15 K to obtain the overall driving force of the pathways and ΔG° of each reaction. Based on the overall driving forces, the nine pathways can be separated into three distinct categories, namely pathways with (a) high (Arginine and Proline Metabolism) (b) medium (Pyruvate, TCA, OXPHOS, PPP, and amino-sugar metabolism) and (c) low (glycolysis/gluconeogenesis and leukotriene metabolism) driving force. The change in the driving force of each pathway is shown in **Figure 1**.

**Figure 1:**
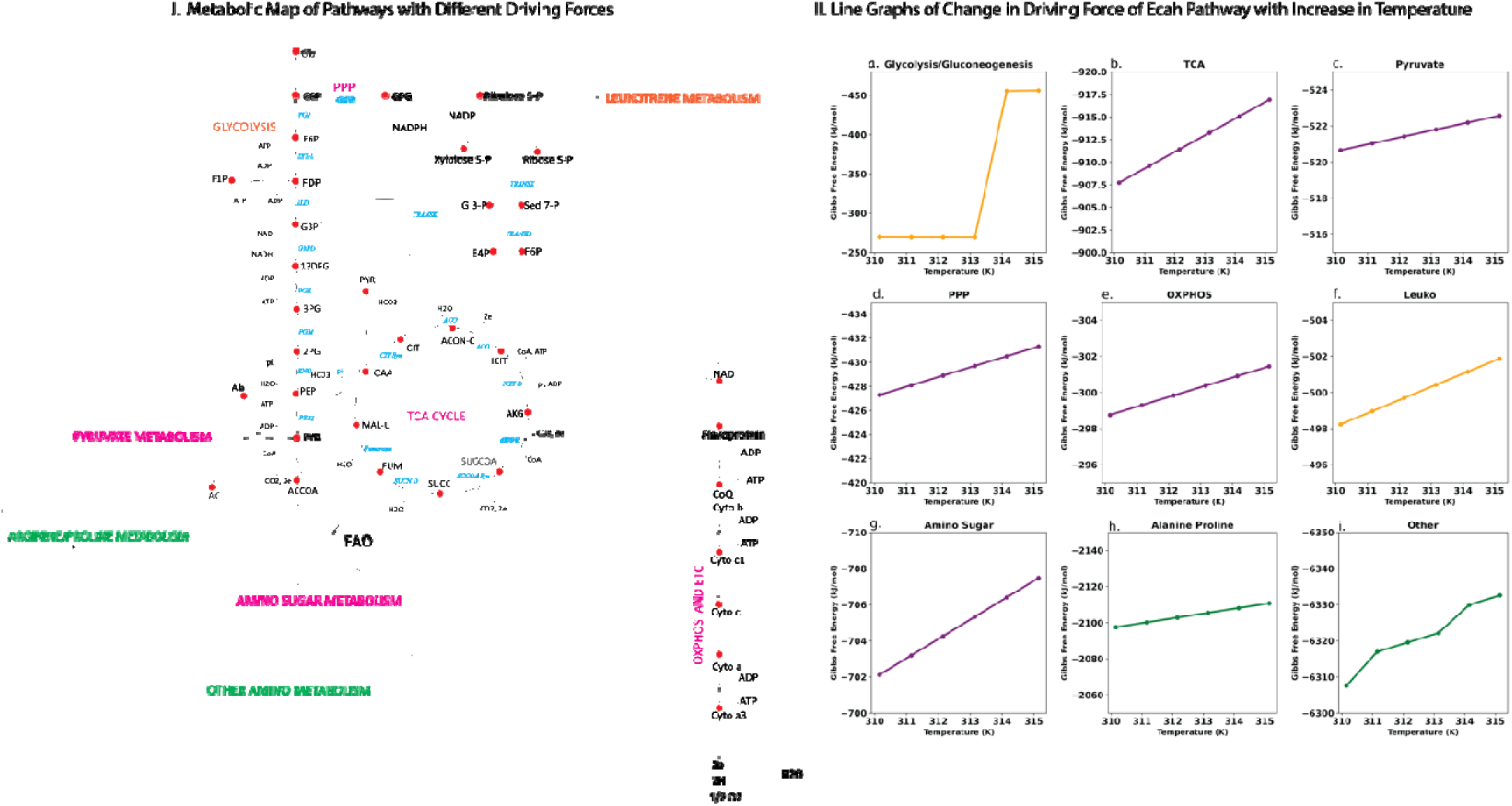
i.) Pathway Map with the pathways with highest driving force highlighted in green, medium driving force highlighted in pink, and lowest driving force highlighted in tan. ii.) The total driving force of the pathways when temperature increases from 310.15 K to 315.15 K for the physiological substrate concentration ranging from 0.01 mM to 10 mM. (a) the change in driving force for the glycolysis/gluconeogenesis pathway which changes from −269.449 to - 456.104 KJ/mol (b) TCA cycle driving force ranging from −907 to −916.9 KJ/mol. (c) Pyruvate metabolism driving force ranging from −520.655 to −522.569 KJ/mol. (d) PPP pathway driving force ranges between −427.285 to −431.306 KJ/mol. (e) OXPHOS driving force range −298.757 to −301.437 KJ/mol. (f) Leukotriene Metabolism driving force range −498.234 to −501.872 KJ/mol. (g) Amino Sugar Metabolism Driving Force −702.114 to −707.474 KJ/mol. (h) Alanine and Proline metabolism driving force: −2097.56 to −2110.77 KJ/mol. (i) Other amino acid metabolism driving force range −6307.54 to −6332.57 KJ/mol.

Although a specific pathway may exhibit a high overall driving force, the individual reactions within it may behave differently from one another. Some reactions can be highly thermodynamically favorable at a certain temperature and that might change with temperature fluctuations. Our MDF predictions are in accordance with the reported thermodynamic nature of pathways. Additionally, the reactions that showed the highest deviation in ΔG° originated mainly from glycolysis/gluconeogenesis and Arginine and Proline metabolism. Glycolytic reactions such as PGI (formation of fructose-1,6-bisphosphate, 3-phospho-D-glycerate), HXK (formation of glucose-6-phosphate), and PYK (the conversion of pyruvate to lactate) were found to have high -ΔG° values from the initial temperature starting at 310.15 K and maintained high values till 315.15 K. The observation that pyruvate production is the most thermodynamically feasible reaction in the glycolytic pathway aligns with findings from several studies^5,8^, reinforcing the reliability of using MDF analysis for further predictions and exploration. Additionally, the reaction glycolaldehyde transferase (formation of sedoheptulose-1,7-bisphosphate) shows a big shift in -ΔG° value around 314.15 K. **Figure 2** shows the shift in thermodynamic feasibility of glycolytic reactions, when temperature increases from 310.15 K to 314.15 K. Similarly, in the Arginine and Proline metabolism, the reaction OAT (use of glutamate for L-glutamate 5-semialdehyde formation) is found to be more thermodynamically infeasible with the temperature increase. However, the formation of L-glutamate 5-semialdehyde via AKG and ornithine consumption was found to increase feasibility between 314.15 and 315.15 K. Similarly, reactions such as 2-oxoglutarate aminotransferase (4-hydroxy-2-oxoglutarate and aspartate formation) start showing an increase in -ΔG° after 3 or more degree rise in temperature. Interestingly, our analysis highlights several metabolites, such as fructose-1,6-bisphosphate, glucose-6-phosphate, and ornithine, which are known^20,29^ to play crucial roles in metabolic regulation and immune responses, as significant contributors.

**Figure 2:**
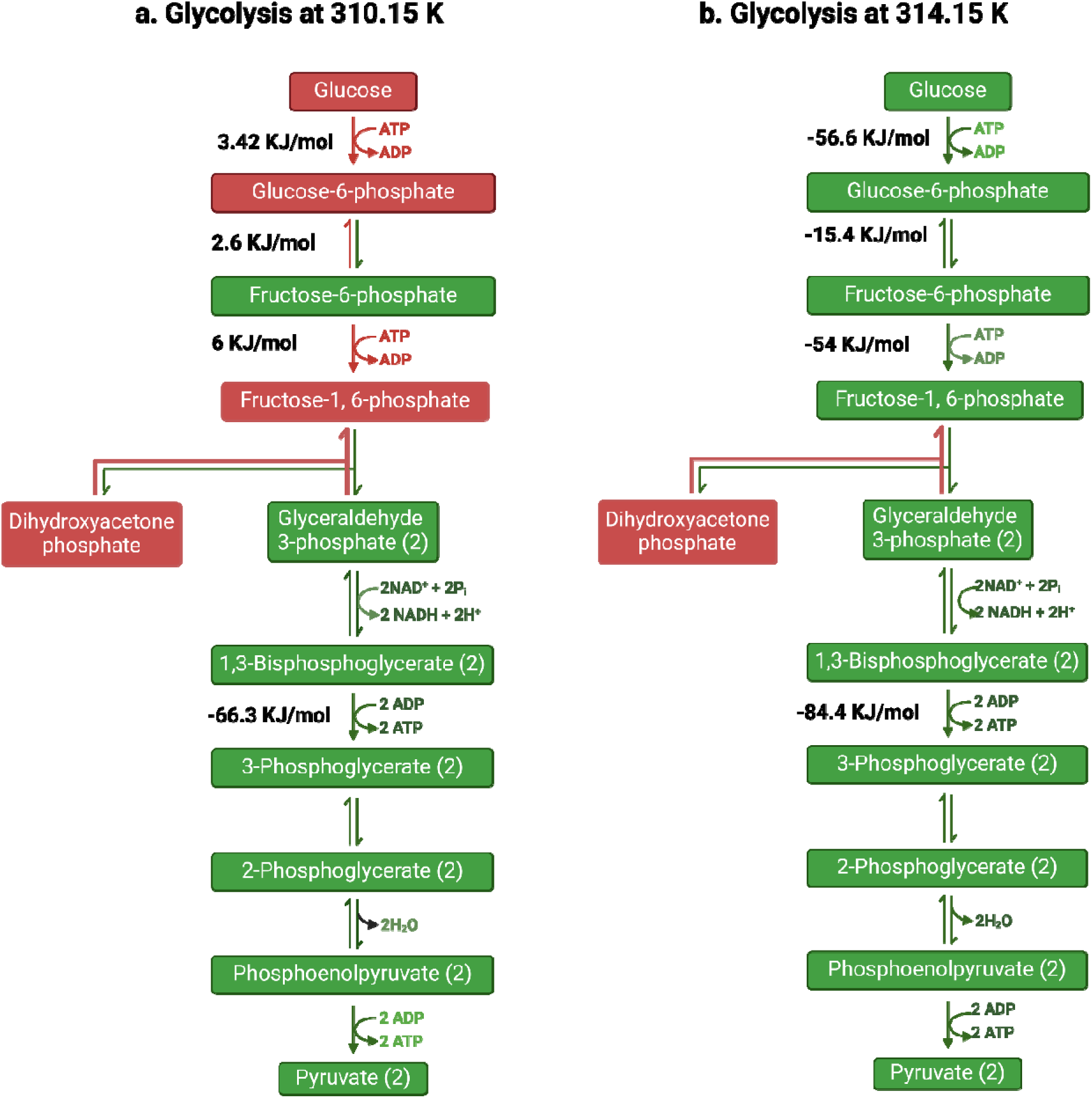
Gibbs free energy of reactions in glycolysis pathway at temperature 310.15 K, which is the normal body temperature, and at 314.15 K where we start to observe dynamic shifts in feasibility of various reactions. a) This pathway map highlights the reactions that have positive Gibbs free energy in red and the reactions with negative Gibbs free energy in green. The red reactions denote thermodynamically infeasible reactions at 310.15 K temperature with green color indicating thermodynamically feasible reactions. These observations are in accordance with the reported phenomena of glycolysis at normal body temperature with the most favorable reaction being the formation of 3-phosphoglycerate from 1,2-biphosphate glycerate and the least favorable reaction being the formation of fructose 1,6-phosphate from DHAP. b) In this metabolic map we highlight the reactions that show a drastic change in Gibbs free energy at 314.15 K temperature. We observed the first three reactions of glycolysis which were first infeasible now to have switched to become highly feasible with an increase in temperature.

**Figure 3:**
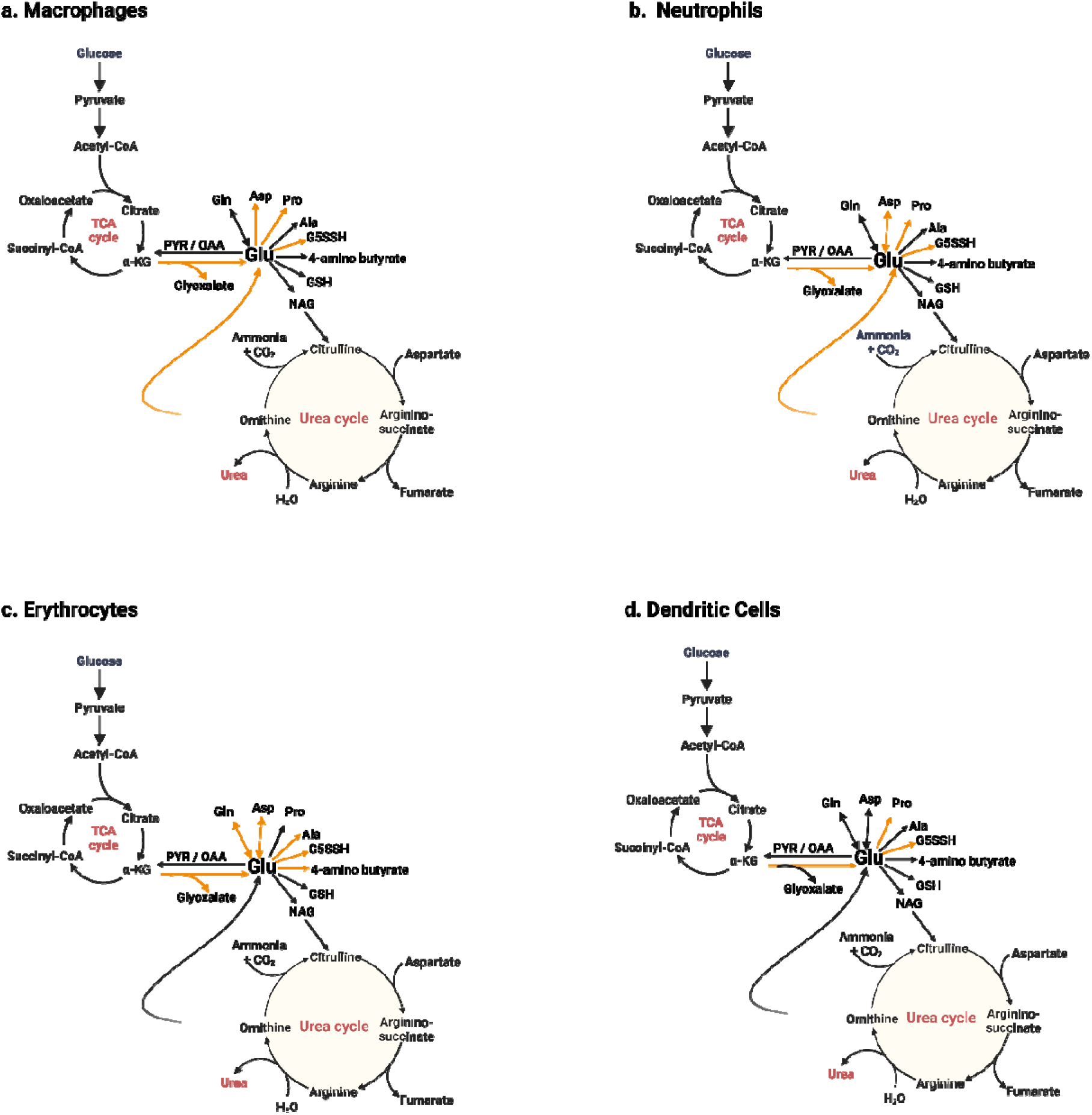
Highlighted reactions that show the highest variation in Gibbs free energy with temperature increase. Orange color denotes the reactions whose Gibbs free energy varies by more than 3KJ/mol with a change in temperature. All the identified reactions for different cell types can be linked to glutamate metabolism a) Metabolic map of reactions associated with glutamate metabolism that show change in Gibbs free energy with temperature increase in LPS induced macrophages. Reactions identified lead to the production of other amino acids such as proline, aspartate, and important compounds like G5SSH. b) Metabolic reactions that are most sensitive to temperature increase in neutrophils which include amino acids such as aspartate, alanine, and proline, and also production of glutamate through ornithine. c) Erythrocyte’s metabolic reactions that show significant Gibbs free energy shift consist of glutamate and glutamine, aspartate, alanine, proline, and 4-amino butyrate. d) Dendritic cells metabolic shift as predicted by Gibbs free energy was found to be limited to mainly proline and G5SSH-associated reactions.

### Change in Reaction Feasibility with Temperature Increase for Immune Cells

We further wanted to explore the change in thermodynamic feasibility of reactions in different immune cells such as macrophages, neutrophils, erythrocytes, and dendritic cells. These immune cells are often the first responders against any disease condition and show highly versatile metabolic behavior. Based on the study by Hooftman et al., 2023^29^, various metabolite concentrations were measured in response to LPS induction in macrophages. Using this data, we adjusted the concentration constraints for metabolites like ATP, ADP, UDP, fumarate, itaconate, and succinate (the complete list of metabolite constraints can be found in the **Supplementary File 1**). We then observed how these adjustments affected the changes in -ΔG° across the nine pathways discussed earlier. LPS induction is the process of exposing macrophages to lipopolysaccharides, which are large molecules found in the outer membrane of gram-negative bacteria^30^. This process acts as a potent activator in macrophages and activates immune responses against bacterial infection^30^. Hence, analyzing the change in feasibility under these conditions allows us to decipher the proper immune response of macrophages against bacterial infection. Similarly, the metabolite concentration trends reported by Kaiser et al., 2020^23^, for neutrophils, erythrocytes, and dendritic cells—specifically metabolites like ornithine, putrescine, and arginine (the complete list is provided in the Supplementary File)—enabled us to perform MDF analysis on four pathways: Arginine/Proline, Alanine, Amino Sugar, and Leukotriene metabolism.

Based on our analysis, the overall category (i.e., high, medium, and low) to which each pathway belongs remained the same with a slight increase or decrease in total driving forces while individual reactions showed unique responses. Numerous reactions involving glutamate seemed to be the most sensitive across different cell types to the temperature increase. To determine what a significant change in ΔG° is, we followed the same criteria as described by Noor et al, 2014^7^ where a change that exceeds approximately 3 KJ/mol indicates a meaningful shift in reaction favorability^7^. **Table S1 in Supplementary File 2** shows the reactions with significant changes in the ΔG° values in macrophages, neutrophils, erythrocytes, and dendritic cells in comparison to the normal state as described earlier. Some of the reactions mentioned earlier, namely PGI and HXK (the formation of fructose1,6-biphosphate, and glucose-6-phosphate) are key metabolites needed for glycolytic switch during the immune response of macrophages^20^. Similarly, numerous metabolites such as pyruvate, various glutamate compounds, and ornithine are important precursors for various amino acid metabolism involved in neutrophil, dendritic cells, and erythrocytes^29^ metabolism. The change in feasibility of the reaction (shown in **Table S1 of Supplementary File 2**) involved in the conversion of arginine to ornithine and urea, conversion of arginine to citrulline, and the formation of agmatine, were found to be more thermodynamically feasible for dendritic and erythrocytes in comparison to neutrophils. The ΔG° values observed across different cell types indicate that the urea cycle plays diverse and cell-specific roles^28,31^. Specifically, this includes the conversion of arginine into ornithine and urea or into citrulline and ammonia. By analyzing shifts in ΔG°, we can gain valuable and reliable insights into the unique metabolic capabilities of each cell type^32^.

Next, we analyzed the reactions with highest change in ΔG° changes with temperature increase within each cell type and found the reactions involving glutamate show distinct increases/decreases. Glutamate metabolism is known to be essential for cellular and physiological functions because of its key roles in various amino acid metabolism, glutamate is a major neurotransmitter in the brain and a precursor for several critical biomolecules^33,34^. To this end, we identified the change in the feasibility of reactions involved with glutamate metabolism as a driving force for each immune cell to acquire its unique function. **Figure 2** highlights the reactions in each cell type that show significant change in their ΔG° value with the temperature fluctuations. We found the fate of the glutamate to be an important factor in determining the function of the cell. For example, in macrophages, the reaction, p5c dehydrogenase, which uses glutamate to produce G5SSH feasibility increases by more than 6 KJ/mol after 2 degrees rise in temperature whereas the reaction, ornithine aminotransferase (OAT), where ornithine and alpha-ketoglutarate produce glutamate and G5SSH show more than 6KJ/mol decrease in ΔG° value after 2-degree rise in temperature. The same reactions, P5C dehydrogenase and OAT, exhibit similar behavior in neutrophils and erythrocytes. However, in dendritic cells, the feasibility of P5C dehydrogenase decreases with an initial rise in temperature and continues to remain below its starting value with each subsequent degree increase. In contrast, the feasibility of the OAT reaction increases with the initial temperature rise and continues to grow steadily up to 315.15 K. Similar analysis reveals the ΔG° value for the reactions involved in proline production increases for all temperature except for the first temperature increase (311.15 K). Overall, how a cell utilizes glutamate to form different products such as ornithine, aspartate, alanine, and 4-amino butyrate (also known as GABA) plays a crucial role in determining the function of a cell type. For example, the change in thermodynamic feasibility of the reaction involved with ornithine in macrophages and neutrophils supports the antimicrobial activity delineated by both cells. Whereas erythrocytes and dendritic cells show more than 6KJ/mol increase in ΔG value for the production of glyoxylate and pyruvate along with a decrease in ΔG° value of reaction-producing aspartate. Hence, we postulate that the fate of glutamate and reprogramming in glutamate metabolism are significant drivers of regulating proper immune responses of different cell types, namely macrophages, neutrophils, dendritic cells, and erythrocytes.

## Discussion

A thermodynamic approach, such as MDF analysis, offers valuable insights into how metabolic and enzymatic pathways in immune cells adapt to thermal fluctuations, including conditions like fever. This method enables the prediction of energy dynamics, reaction sensitivities, and oxidative stress management, enhancing our understanding of immune responses and identifying potential therapeutic targets. In this study, we applied MDF analysis to investigate the impact of temperature increases on nine distinct pathways and examined the thermodynamic feasibility of these pathways across different immune cell types.

Temperature plays a key role in the thermodynamics of metabolic reactions that are critical for maintaining physiological balance, optimizing metabolic engineering processes, and informing clinical practices^1^. ΔG° is fundamental for assessing reaction feasibility in various biological and biotechnological contexts as it quantifies the spontaneity of a reaction^4^. Hence by using MDF analysis, we can quantify the spontaneity of reactions at different temperatures and metabolite concentrations. However, due to limitations on the availability of standard Gibbs free energy values, we were able to analyze eight full pathways and one compilation of amino acid/ fatty acid reactions for which standard Gibbs free energy values can be estimated through a tool called equilibrator^14^. The pathways in this study (such as glycolysis, TCA, OXPHOS, amino acid metabolisms, etc) are crucial for energy metabolism and immune response regulation. While including a greater number of pathways and reactions would provide valuable insights, the nine pathways we have included in this study can provide insights into key metabolic capabilities and immune responses at different temperatures. We started our analysis with a normal body temperature of 310.15 K and went up to 315.15 K, which is known to be deleterious to health.

One of the challenges in conducting such thermodynamic analysis is to be able to gather proper concentration data. In this study, we used a range of 0.01 mM to 10 mM for normal conditions and to be able to capture the immune cell phenotypes we used the reported metabolite concentration trends reported by Hooftman et al, 2023^21^ for LPS-induced macrophages and Kaiser et al, 2020^23^ for neutrophil, dendritic cells, and erythrocytes. While we were not able to use specific concentrations for all the metabolites involved in the nine pathways mentioned above, the list of metabolites we obtained from these two studies are highly significant metabolites such as ATP, ADP, and various amino acids such as arginine, glutamate, and proline that play crucial roles in metabolism and immune response. Changing the concentration of these metabolites showed a significant impact on the feasibility of the pathways and the associated reactions, which allowed us to garner insights into how the metabolic shift occurs with the temperature increase. However, further advancements in tools for standard Gibbs free energy prediction and metabolite concentrations for different cell-types will enable more comprehensive thermodynamic analysis. This will enable the proper study of complex immune responses, which will allow the identification of effective and novel therapeutic targets.

Through MDF analysis, we found that the overall feasibility of all pathways increased with temperature however, reaction feasibility differed highlighting reactions from glycolysis/gluconeogenesis, Arginine/Proline, Pyruvate, and PPP that could be significant for normal physiological activity and for immune responses. In doing so, we identified glutamate compound derivatives namely glutamate 5-semialdehyde and ornithine to show significant change in thermodynamic feasibility across the mentioned cell types. Glutamate-5-semialdehyde is an important intermediate in the biosynthesis of proline, which ultimately contributes to the formation of GABA. In human immune cells, the conversion of glutamate to glutamate-5-semialdehyde, and subsequently to ornithine and other amino acids, plays a significant role in regulating metabolic pathways that support immune responses. These pathways are particularly relevant in the context of cellular stress, where proline and GABA may contribute to modulating oxidative stress and maintaining immune cell function, drawing parallels to the roles observed in plant cells under abiotic stress conditions^20^.

Similarly, Ginguay et al.^32,33^ investigated the role of glutamate and its derivatives in modulating the human immune response and regulating fate determination. However, the specific utilization of glutamate to form different compounds within immune cells has not been as exhaustively explored^35^. Glutamate, derived from glutamine through glutaminolysis, plays roles in amino acid synthesis and as a neurotransmitter. In immune cells, glutamate can be further metabolized into compounds such as α-ketoglutarate, which enters the tricarboxylic acid (TCA) cycle, influencing energy production and biosynthesis^36^. Additionally, glutamate serves as a precursor for the synthesis of glutathione, a vital antioxidant that protects immune cells from oxidative stress^37^. Emerging research highlights the importance of glutamate metabolism in immune function. For instance, the conversion of glutamate-derived metabolites like fumarate has been linked to epigenetic reprogramming in monocytes, enhancing their pathogen responsiveness, a process termed “trained immunity”^37^. While glutamine metabolism is well-characterized in immune responses, the specific roles of glutamate in supporting cellular functions in macrophages, neutrophils, dendritic cells, and erythrocytes are only beginning to be understood. Through thermodynamic analysis of reactions and pathways, we emphasize the potential significance of glutamate metabolism in immune cell functionality. **Table S1 in Supplementary File 2** provides a detailed overview of reactions showing distinct shifts in Gibbs free energy, underlining the energetic and regulatory importance of glutamate in modulating immune responses.

## Conclusion

MDF analysis provides reliable insight into pathway thermodynamics and reaction feasibility and accurately captures the shift due to temperature and metabolite concentrations. We were able to categorize the pathways into three distinct categories and highlight the reactions that show dynamic shifts with temperature increase. We also postulate the fate of glutamate to be one of the most important factors in modulating immune responses for different immune cells. Through this study, we aim to highlight the potential of using thermodynamic analysis such as MDF in navigating the complex human metabolism and activation of proper immune responses.

## Method

### Standard Gibbs Free Energy and Max/Min Driving Force

Δ*_f_*G° was calculated for reactions using the equilibrator tool^20^. Equilibrator is a tool that uses the composition contribution method to calculate the Gibbs free energy of formation at standard conditions. The list of reactions and the obtained standard Δ*_f_*G° values are available in the **Supplementary File 1**. After acquiring the list of standard Gibbs Free Energy, MDF) or the pathways of interest were calculated by using the concentration of metabolites ranging from 0.01 mM to 10 mM, which is the normal physiological range of metabolites maintained in a human cell^38^. The details on the calculation of MDF and change in ΔG° can be found in Elad Noor et al, 2010^7^, and Chowdhury et al, 2023^26^.

## Abbreviations

MDF: Max/Min Driving Force
Δ*_f_*G°: standard Gibbs free energy
ΔG°: Gibbs free energy
OXPHOS: Oxidative Phosphorylation
TCA: Tricarboxylic Acid Cycle
PPP: Pentose Phosphate Pathway
PGI: Phosphofructokinase
HXK: Hexokinase
PYK: Pyruvate Kinase
OAT: Ornithine aminotransferase
AKG: Alpha ketoglutarate
GABA: Gamma-Aminobutyric Acid
ATP: Adenosine Triphosphate
ADP: Adenosine Diphosphate
UDP: Uridine Diphosphate
CCM: Component Contribution Method
G5SSH: L-glutamate 5-semialdehyde
TMFA: Metabolic Flux Analysis

## Declarations

### Ethics approval and consent to participate

N/A

### Consent for publication

N/A

### Availability of data and material

All data generated or analyzed during this study are included in this published article and its supplementary information files. The code for the MDF analysis used in this study is available in this GitHub repository: https://github.com/ChowdhuryRatul/kcat_iZMA6517.

### Authors’ contributions

S.M. and R.S. worked on concept development for this work and developed the methodologies. The analysis followed by the writing was done by S.M. and R.S.

## Acknowledgments

We gratefully acknowledge the Holland Computing Center (HCC) of the University of Nebraska, which receives support from the Nebraska Research Initiative (United States of America).

## References

1. Evans, S. S., Repasky, E. A. & Fisher, D. T. Fever and the thermal regulation of immunity: the immune system feels the heat. Nat. Rev. Immunol. 15, 335–349 (2015).

2. Wang, H., Ülgen, M. & Trajkovski, M. Importance of temperature on immuno-metabolic regulation and cancer progression. FEBS J. 291, 832–845 (2024).

3. Deciphering the relationship between temperature and immunity | Discovery Immunology | Oxford Academic. https://academic.oup.com/discovimmunology/article/3/1/kyae001/7591951.

4. Noor, E., Haraldsdóttir, HS., Milo, R. & Fleming, RMT. Consistent Estimation of Gibbs Energy Using Component Contributions. PLOS Comput. Biol. 9, e1003098 (2013).

5. Greinert, T., Vogel, K., Maskow, T. & Held, C. New thermodynamic activity-based approach allows predicting the feasibility of glycolysis. Sci. Rep. 11, 6125 (2021).

6. Entropy-favored human antibody binding reactions with a non-infectious antigen - ScienceDirect. https://www.sciencedirect.com/science/article/pii/S0161589007007389?via%3Dihub.

7. Noor, E. et al. Pathway Thermodynamics Highlights Kinetic Obstacles in Central Metabolism. PLOS Comput. Biol. 10, e1003483 (2014).

8. Henry, C. S., Broadbelt, L. J. & Hatzimanikatis, V. Thermodynamics-Based Metabolic Flux Analysis. Biophys. J. 92, 1792–1805 (2007).

9. Ataman, M. & Hatzimanikatis, V. Heading in the right direction: thermodynamics-based network analysis and pathway engineering. Curr. Opin. Biotechnol. 36, 176–182 (2015).

10. Neutrophils Fuel Effective Immune Responses through Gluconeogenesis and Glycogenesis: Cell Metabolism. https://www-cell-com.libproxy.unl.edu/cell-metabolism/fulltext/S1550-4131(20)30651-3.

11. Remmerie, A. & Scott, C. L. Macrophages and lipid metabolism. Cell. Immunol. 330, 27–42 (2018).

12. Møller, S. H., Wang, L. & Ho, P.-C. Metabolic programming in dendritic cells tailors immune responses and homeostasis. Cell. Mol. Immunol. 19, 370–383 (2022).

13. Hess, J. R., Rugg, N., Joiner, C. H., Jacobs, M. R. & Klein, H. G. Red blood cell metabolism under prolonged anaerobic storage. Mol. BioSyst. 9, 2555–2575 (2013).

14. NIST. Data. National Institute of Standards and Technology (2016).

15. Kanehisa, M., Furumichi, M., Sato, Y., Kawashima, M. & Ishiguro-Watanabe, M. KEGG for taxonomy-based analysis of pathways and genomes. Nucleic Acids Res. 51, D587–D592 (2023).

16. MetaNetX/MNXref: unified namespace for metabolites and biochemical reactions in the context of metabolic models | Nucleic Acids Research | Oxford Academic. https://academic-oup-com.libproxy.unl.edu/nar/article/49/D1/D570/5958493.

17. Norsigian, C. J. et al. BiGG Models 2020: multi-strain genome-scale models and expansion across the phylogenetic tree. Nucleic Acids Res. 48, D402–D406 (2020).

18. ChEBI in 2016: Improved services and an expanding collection of metabolites | Nucleic Acids Research | Oxford Academic. https://academic-oup-com.libproxy.unl.edu/nar/article/44/D1/D1214/2502583.

19. Henry, C. S. et al. High-throughput generation, optimization and analysis of genome-scale metabolic models. Nat. Biotechnol. 28, 977–982 (2010).

20. Beber, M. E. et al. eQuilibrator 3.0: a database solution for thermodynamic constant estimation. Nucleic Acids Res. 50, D603–D609 (2022).

21. Hubbard, R. W., Bowers, W. D., Matthew, W. T. & Curtis, F. C. Alteration in circulating metabolites during and after heat stress in the conscious rat: potential biomarkers of exposure and organ-specific injury. BMC Physiol. 14, 14 (2014).

22. Schranner, D., Kastenmüller, G., Schönfelder, M., Römisch-Margl, W. & Wackerhage, H. Metabolite Concentration Changes in Humans After a Bout of Exercise: a Systematic Review of Exercise Metabolomics Studies. Sports Med. - Open 6, 11 (2020).

23. Kaiser, L. et al. Metabolite Patterns in Human Myeloid Hematopoiesis Result from Lineage-Dependent Active Metabolic Pathways. Int. J. Mol. Sci. 21, 6092 (2020).

24. Watanabe, I. & Okada, S. EFFECTS OF TEMPERATURE ON GROWTH RATE OF CULTURED MAMMALIAN CELLS (L5178Y). J. Cell Biol. 32, 309–323 (1967).

25. Summary, conclusions and recommendations: adverse temperature levels in the human body: International Journal of Hyperthermia: Vol 19, No 3. https://www-tandfonline-com.libproxy.unl.edu/doi/abs/10.1080/0265673031000090701.

26. Chowdhury, N. B. et al. A multi-organ maize metabolic model connects temperature stress with energy production and reducing power generation. iScience 26, 108400 (2023).

27. Goldberg, R. N. & Tewari, Y. B. Thermodynamics of the hydrolysis reactions of adenosine 3′,5′-(cyclic)phosphate(aq) and phospho*enol*pyruvate(aq); the standard molar formation properties of 3′,5′-(cyclic)phosphate(aq) and phospho*enol*pyruvate(aq). J. Chem. Thermodyn. 35, 1809–1830 (2003).

28. Steinhauser, M. L., et al. The circulating metabolome of human starvation. JCI Insight 3, (2018).

29. Macrophage fumarate hydratase restrains mtRNA-mediated interferon production | Nature. https://www-nature-com.libproxy.unl.edu/articles/s41586-023-05720-6#Sec43.

30. Meng, F. & Lowell, C. A. Lipopolysaccharide (LPS)-induced Macrophage Activation and Signal Transduction in the Absence of Src-Family Kinases Hck, Fgr, and Lyn. J. Exp. Med. 185, 1661 (1997).

31. Martí i Líndez, A.-A. & Reith, W. Arginine-dependent immune responses. Cell. Mol. Life Sci. CMLS 78, 5303–5324 (2021).

32. Ganeshan, K. & Chawla, A. Metabolic Regulation of Immune Responses. Annu. Rev. Immunol. 32, 609–634 (2014).

33. Ginguay, A., Cynober, L., Curis, E. & Nicolis, I. Ornithine Aminotransferase, an Important Glutamate-Metabolizing Enzyme at the Crossroads of Multiple Metabolic Pathways. Biology 6, 18 (2017).

34. Hansen, A. M. & Caspi, R. R. Glutamate joins the ranks of immunomodulators. Nat. Med. 16, 856 (2010).

35. Brosnan, J. T. Glutamate, at the Interface between Amino Acid and Carbohydrate Metabolism. J. Nutr. 130, 988S–990S (2000).

36. Tapiero, H., Mathé, G., Couvreur, P. & Tew, K. D. II. Glutamine and glutamate. Biomed. Pharmacother. 56, 446–457 (2002).

37. Ganor, Y. & Levite, M. Glutamate in the Immune System: Glutamate Receptors in Immune Cells, Potent Effects, Endogenous Production and Involvement in Disease. in Nerve-Driven Immunity: Neurotransmitters and Neuropeptides in the Immune System (ed. Levite, M.) 121–161 (Springer, Vienna, 2012). doi:10.1007/978-3-7091-0888-8_4.

38. Substrate Concentration - an overview | ScienceDirect Topics. https://www.sciencedirect.com/topics/nursing-and-health-professions/substrate-concentration.

