## Supplementary Table 1 for "Pathway Thermodynamic Analysis Postulates Change in Glutamate Metabolism as a Key Factor in Modulating Immune Responses"

**Supplementary Table for the study “Pathway Thermodynamic Analysis Identifies Change in Glutamate Metabolism as a Key Factor in Modulating Immune Responses”**

**Table S1:** The table below shows the complete list of reactions originating from the analyzed pathways that show distinct shift in thermodynamic feasibility for different cell types. The table consist of four columns where the first column is the reaction definition, followed by the Pathway. The third column shows the MDF prediction on how the shift on Gibbs free energy occurs which is the further explained in column 4 with possible contribution of said change. Finally, the last column mentioned which cell type the change occurs in.

| Reaction | Pathway | Change Observed | Outcome | Cell type |
| --- | --- | --- | --- | --- |
| fructose-6-phosphate[c] + UTP[c] => fructose-1,6-bisphosphate[c] + H+[c] + UDP[c] | Glycolysis/gluconeogenesis | +∆G^o^ to - ∆G^o^ | Enables switch to glycolytic metabolism^1^ | Macrophage |
| 1,3-bisphospho-D-glycerate[c] + ADP[c] <=> 3-phospho-D-glycerate[c] + ATP[c] | Glycolysis/gluconeogenesis | more than 4X change in ∆G^o^ value | Precursors for amino acids such as serine, cysteine, glycine, and alanine are crucial for essential cellular functions^2,3^. | Macrophage |
| 1,3-bisphospho-D-glycerate[c] + H2O[c] => 3-phospho-D-glycerate[c] + H+[c] + Pi[c] | Glycolysis/gluconeogenesis | More than 10KJ/mol change in ∆G^o^ value | Precursors for amino acids such as serine, cysteine, glycine, and alanine are important for essential cellular functions^2,3^. | Macrophage |
| acetoacetate[m] + succinyl-CoA[m] => acetoacetyl-CoA[m] + succinate[m] | TCA | +∆G^o^ to - ∆G^o^ | Helps in the generation of the mitochondrial proton gradient and ATP synthesis^4^ | Macrophage |
| isocitrate[m] + NAD+[m] => AKG[m] + CO2[m] + NADH[m] | TCA | +∆G^o^ to - ∆G^o^ | Helps in the generation of the mitochondrial proton gradient and ATP synthesis^4^ | Macrophage |
| glyoxalate[m] + H+[m] + NADPH[m] => glycolate[m] + NADP+[m] | TCA | +∆G^o^ to - ∆G^o^ | Suppresses harmful effects of peroxide on mitochondrial activity^5^. | Macrophage |
| H+[c] + NADH[c] + OAA[c] <=> malate[c] + NAD+[c] | TCA | More than 10KJ/mol change in ∆G^o^ value | Helps in the generation of the mitochondrial proton gradient and ATP synthesis^4^ | Macrophage |
| CoA[m] + GTP[m] + succinate[m] <=> GDP[m] + Pi[m] + succinyl-CoA[m] | TCA | Decrease in ∆G^o^ value | Helps in the generation of the mitochondrial proton gradient and ATP synthesis^4^ | Macrophage |
| glycolaldehyde[c] + H2O[c] + NAD+[c] => glycolate[c] + 2 H+[c] + NADH[c] | TCA | Decrease in ∆G^o^ value | Suppresses harmful effects of peroxide on mitochondrial activity^5^. | Macrophage |
| glycolaldehyde[m] + H2O[m] + NAD+[m] => glycolate[m] + 2 H+[m] + NADH[m] | TCA | More than 10KJ/mol change in ∆G^o^ value | Suppresses harmful effects of peroxide on mitochondrial activity^5^. | Macrophage |
| H+[c] + lactaldehyde[c] + NADPH[c] => NADP+[c] + propane-1,2-diol[c] | Pyruvate Metabolism | +∆G^o^ to - ∆G^o^ | Alterations in carbohydrate metabolism^6^. | Macrophage |
| malate[m] + NAD+[m] => CO2[m] + NADH[m] + pyruvate[m] | Pyruvate Metabolism | +∆G^o^ to - ∆G^o^ | Precursors for amino acids such as serine, cysteine, glycine, and alanine are very important for essential cellular functions^2,3^. | Macrophage |
| H+[m] + NADH[m] + pyruvate[m] <=> L-lactate[m] + NAD+[m] | Pyruvate Metabolism | -∆G^o^ to + ∆G^o^ | Helps in up-regulation of glycolysis^1^. | Macrophage |
| malate[m] + NADP+[m] => CO2[m] + NADPH[m] + pyruvate[m] | Pyruvate Metabolism | More than 10KJ/mol change in ∆G^o^ value | Precursors for amino acids such as serine, cysteine, glycine, and alanine are very important for essential cellular functions^2,3^. | Macrophage |
| acetyl-CoA[c] + H2O[c] => acetate[c] + CoA[c] + H+[c] | Pyruvate Metabolism | More than 10KJ/mol change in ∆G^o^ value | Promotes anti-tumor activity in immune cells^7^. | Macrophage |
| 2-deoxy-D-ribose-5-phosphate[c] <=> acetaldehyde[c] + GAP[c] | PPP | +∆G^o^ to - ∆G^o^ | Inflammatory mediator in immune cells^8^. | Macrophage |
| ATP[c] + deoxyribose[c] => 2-deoxy-D-ribose-5-phosphate[c] + ADP[c] + H+[c] | PPP | +∆G^o^ to - ∆G^o^ | Changes in mitochondrial function, possibly provoking of oxidative stress^9^. | Macrophage |
| 5 H+[m] + NADH[m] + ubiquinone[m] => NAD+[m] + ubiquinol[m] + 4 H+[i] | OXPHOS | More than 10KJ/mol change in ∆G^o^ value | Promotes mitochondrial function^1^. | Macrophage |
| O2[r] + AKG[r] + proline[r] => CO2[r] + succinate[r] + trans-4-hydroxy-L-proline[r] | Alanine Proline Metabolism | More than 10KJ/mol change in ∆G^o^ value | Precursors for amino acids such as serine, cysteine, glycine, and alanine are very important for essential cellular functions^2,3^. | Macrophage |
| arginine[c] + 2 NADPH[c] + 2 O2[c] => citrulline[c] + 2 H2O[c] + 2 NADP+[c] + NO[c] | Alanine Proline Metabolism | More than 10KJ/mol change in ∆G^o^ value | Regulation of urea cycle for proper immune response^10^. | Macrophage |
| ATP[c] + glucosamine[c] => ADP[c] + glucosamine-6-phosphate[c] + H+[c] | Amino sugar metabolism | More than 10KJ/mol change in ∆G^o^ value | Inhibition of inflammation^11^. | Macrophage |
| ATP[c] + glucosamine[c] => ADP[c] + glucosamine-6-phosphate[c] + H+[c] | Amino sugar metabolism | More than 10KJ/mol change in ∆G^o^ value | Inhibition of inflammation^11^. | Macrophage |
| glutamate[m] + 2 H+[m] + NADH[m] <=> H2O[m] + L-glutamate 5-semialdehyde[m] + NAD+[m] | Arginine and proline metabolism | More than 6 KJ/mol increase | Precursor to other amino acids, involvement in energy metabolism, and immune cell activation^25^. | Neutrophils, Dendritic cells, |
| AKG[m] + ornithine[m] <=> glutamate[m] + L-glutamate 5-semialdehyde[m] | Arginine and proline metabolism | More than 5 KJ/mol increase | The precursor to polyamine synthesis helps in the regulation of NO cycle^30,31^. | Neutrophil; |
| arginine[c] + H2O[c] => citrulline[c] + H+[c] + NH3[c] | Arginine and proline metabolism | More than 6 KJ/mol increase | Helps in the regulation of the NO cycle and promotes the antimicrobial activity^30^. | Neutrophil |
| glutamyl-5-phosphate[m] + H+[m] + NADPH[m] => L-glutamate 5-semialdehyde[m] + NADP+[m] + Pi[m] | Arginine and proline metabolism | More than 6 KJ/mol increase | Precursor to other amino acids, involvement in energy metabolism, and immune cell activation^25^. | Neutrophil |
| arginine[m] + H+[m] => agmatine[m] + CO2[m] | Arginine and proline metabolism | More than 6 KJ/mol increase | The precursor to polyamines helps in immune modulation^31,32^. | Neutrophil |
| AKG[m] + L-erythro-4-hydroxyglutamate[m] => 4-hydroxy-2-oxoglutarate[m] + glutamate[m] | Arginine and proline metabolism | More than 6 KJ/mol decrease | Precursor to other amino acids, involvement in energy metabolism, and immune cell activation^25^. | Neutrophil |
| acetyl-CoA[m] + glutamate[m] => CoA[m] + H+[m] + N-acetyl-L-glutamate[m] | Arginine and proline metabolism | More than 6 KJ/mol decrease | The precursor to other amino acids essential for cellular function^12,13^. | Neutrophil |
| arginine[m] + H2O[m] => ornithine[m] + urea[m] | Arginine and proline metabolism | More than 7 KJ/mol increase | Balance of NO cycle and maintain proper inflammatory response^14^. | Neutrophil |
| O2[r] + AKG[r] + proline[r] => CO2[r] + succinate[r] + trans-4-hydroxy-L-proline[r] | Arginine and proline metabolism | More than 5 KJ/mol decrease | Precursor to other amino acids, involvement in energy metabolism, and immune cell activation^25^. | Neutrophil |
| arginine[c] + 2 NADPH[c] + 2 O2[c] => citrulline[c] + 2 H2O[c] + 2 NADP+[c] + NO[c] | Arginine and proline metabolism | More than 6 KJ/mol decrease | Balance of NO cycle and maintain proper inflammatory response^15^. | Neutrophil |
| serine[c] => H+[c] + NH3[c] + pyruvate[c] | Glycine, serine and threonine metabolism | More than 5 KJ/mol decrease | Maintains cellular function and acts as a precursor for synthesizing proteins, nucleotides, and other biomolecules^1^. | Neutrophil |
| serine[c] => dehydroalanine[c] + H2O[c] | Glycine, serine and threonine metabolism | More than 4 KJ/mol decrease | Maintains cellular function and acts as a precursor for synthesizing proteins, nucleotides, and other biomolecules^1^. | Neutrophil |
| arginine[c] + glycine[c] <=> guanidinoacetate[c] + ornithine[c] | Glycine, serine and threonine metabolism | More than 5 KJ/mol decrease | The precursor to polyamine synthesis helps in the regulation of NO cycle^14^. | Neutrophil |
| alanine[m] + glyoxalate[m] => glycine[m] + pyruvate[m] | Glycine, serine and threonine metabolism | More than 6 KJ/mol decrease | Essential for amino acid metabolism^1^. | Neutrophil |
| alanine[x] + glyoxalate[x] => glycine[x] + pyruvate[x] | Glycine, serine and threonine metabolism | More than 5 KJ/mol decrease | Essential for amino acid metabolism^21^. | Neutrophil |
| aspartate[c] + ATP[c] + citrulline[c] => AMP[c] + argininosuccinate[c] + H+[c] + PPi[c] | Alanine, aspartate and glutamate metabolism | More than 5 KJ/mol decrease | A crucial step in incorporating nitrogen into the urea cycle^10,15^. | Neutrophil |
| aspartate[c] => fumarate[c] + H+[c] + NH3[c] | Alanine, aspartate and glutamate metabolism | More than 6 KJ/mol decrease | A crucial step for the deamination of amino acids to aid energy metabolism^16^. | Neutrophil |
| ATP[c] + glutamate[c] + NH3[c] => ADP[c] + glutamine[c] + Pi[c] | Alanine, aspartate and glutamate metabolism | More than 4 KJ/mol decrease | Essential for maintaining nitrogen balance and detoxifying ammonia^12,17^. | Neutrophil |
| AKG[c] + alanine[c] <=> glutamate[c] + pyruvate[c] | Alanine, aspartate and glutamate metabolism | More than 5 KJ/mol increase | Essential for maintaining nitrogen balance and detoxifying ammonia^12^. | Neutrophil |
| AKG[m] + alanine[m] <=> glutamate[m] + pyruvate[m] | Alanine, aspartate and glutamate metabolism | More than 8 KJ/mol decrease | Essential for maintaining nitrogen balance and detoxifying ammonia^23^. | Neutrophil |
| 2-oxoglutaramate[c] + H2O[c] => AKG[c] + H+[c] + NH3[c] | Alanine, aspartate and glutamate metabolism | More than 5 KJ/mol decrease | Helps to manage proper ammonia levels and NO^14^. | Neutrophil |
| asparagine[c] + H2O[c] => aspartate[c] + H+[c] + NH3[c] | Alanine, aspartate and glutamate metabolism | More than 7 KJ/mol increase | Helps to manage proper ammonia level^14^. | Neutrophil |
| glutamine[c] + pyruvate[c] => 2-oxoglutaramate[c] + alanine[c] | Alanine, aspartate and glutamate metabolism | -∆G^o^ to +∆G^o^ | Essential for maintaining nitrogen balance and detoxifying ammonia^12,18^. | Neutrophil |
| glutamine[m] + pyruvate[m] => 2-oxoglutaramate[m] + alanine[m] | Alanine, aspartate and glutamate metabolism | +∆G^o^ to _-∆G^o^ | Essential for maintaining nitrogen balance and detoxifying ammonia^22,23^. | Neutrophil |
| D-alanine[c] <=> alanine[c] | Alanine, aspartate and glutamate metabolism | 0 to - ∆G^o^ |  | Neutrophil |
| H2O[c] + N-acetyl-L-aspartate[c] => acetate[c] + aspartate[c] | Alanine, aspartate and glutamate metabolism | More than 4 KJ/mol increase | Helps to manage proper acetate supply^7^. | Neutrophil |
| 1-pyrroline-5-carboxylate[c] + H+[c] + H2O[c] <=> L-glutamate 5-semialdehyde[c] | Arginine and proline metabolism | More than 4 KJ/mol decrease | Precursor to other amino acids, involvement in energy metabolism, and immune cell activation^25^. | Dendritic cells |
| CO2[m] + H2O[m] => H+[m] + HCO3-[m] | Arginine and proline metabolism | More than 6 KJ/mol increase | Important for ph regulation and Co2 transport for metabolic processes^19^. | Dendritic cells |
| 5-methylthioadenosine[c] + Pi[c] => adenine[c] + methylthioribose-1p[c] | Arginine and proline metabolism | More than 4 KJ/mol decrease | Precursor for methionine and nucleotides^20^. | Dendritic cells |
| acetyl-CoA[m] + glutamate[m] => CoA[m] + H+[m] + N-acetyl-L-glutamate[m] | Arginine and proline metabolism | More than 5 KJ/mol increase | Precursor to other amino acids essential for cellular function. | Dendritic cells |
| H2O[c] + N-acetylornithine[c] <=> acetate[c] + ornithine[c] | Arginine and proline metabolism | More than 5 KJ/mol increase | Regulation of NO cycle and antimicrobial activity^14^. | Dendritic cells |
| acetyl-CoA[c] + putrescine[c] => CoA[c] + H+[c] + N-acetylputrescine[c] | Arginine and proline metabolism | More than 4 KJ/mol decrease | Balance and proper functioning of polyamines^21^. | Dendritic cells |
| AKG[c] + L-erythro-4-hydroxyglutamate[c] => glutamate[c] + 4-hydroxy-2-oxoglutarate[c] | Arginine and proline metabolism | More than 3 KJ/mol decrease | Maintenance of amino acid and nitrogen metabolism^37^. | Dendritic cells |
| serine[c] + THF[c] <=> 5,10-methylene-THF[c] + glycine[c] + H2O[c] | Glycine, serine and threonine metabolism | More than 3 KJ/mol increase | Maintains cellular function and acts as a precursor for synthesizing proteins, nucleotides, and other biomolecules^20^. | Dendritic cells |
| CoA[m] + L-2-amino-3-oxobutanoic acid[m] <=> acetyl-CoA[m] + glycine[m] | Glycine, serine and threonine metabolism | More than 3 KJ/mol decrease | Maintains cellular function and acts as a precursor for synthesizing proteins, nucleotides, and other biomolecules^42^. | Dendritic cells |
| dehydroalanine[c] + H2O[c] => H+[c] + NH3[c] + pyruvate[c] | Glycine, serine and threonine metabolism | More than 3 KJ/mol decrease | Precursor for other amino acid metabolism^37^. | Dendritic cells |
| serine[c] => dehydroalanine[c] + H2O[c] | Glycine, serine and threonine metabolism | More than 3 KJ/mol increase of + ∆G^o^ | Maintains cellular function and acts as a precursor for synthesizing proteins, nucleotides, and other biomolecules^42^. | Dendritic cells |
| 1-pyrroline-5-carboxylate[m] + 2 H2O[m] + NAD+[m] => glutamate[m] + H+[m] + NADH[m] | Alanine, aspartate and glutamate metabolism | More than 3 KJ/mol decrease | Maintains cellular function and acts as a precursor for synthesizing proteins, nucleotides, and other biomolecules^42^. | Dendritic cells |
| arginine[c] + H2O[c] => ornithine[c] + urea[c] | Arginine and proline metabolism | More than 3 KJ/mol decrease | Balance of NO cycle and ammonia levels^37^. | Erythrocytes |
| H+[e] + ornithine[e] => CO2[e] + putrescine[e] | Arginine and proline metabolism | More than 5 KJ/mol increase | Balance of NO cycle and ammonia levels^14^. | Erythrocytes |
| AKG[m] + L-erythro-4-hydroxyglutamate[m] => 4-hydroxy-2-oxoglutarate[m] + glutamate[m] | Arginine and proline metabolism | More than 5 KJ/mol increase | Precursor to other amino acids, involvement in energy metabolism, and immune cell activation^25^. | Erythrocytes |
| 4-hydroxy-2-oxoglutarate[m] => glyoxalate[m] + pyruvate[m] | Arginine and proline metabolism | More than 5 KJ/mol decrease | Precursor for many important amino acid metabolism^17^. | Erythrocytes |
| H2O[c] + N1-acetylspermidine[c] + O2[c] => acetamidopropanal[c] + H2O2[c] + putrescine[c] | Arginine and proline metabolism | More than 3 KJ/mol decrease | Regulation of cellular growth, and maintenance of proper level of H2O2^22^. | Erythrocytes |
| L-erythro-4-hydroxyglutamate[m] + OAA[m] => 4-hydroxy-2-oxoglutarate[m] + aspartate[m]  arginine[m] + H2O[m] => ornithine[m] + urea[m] | Arginine and proline metabolism | More than 4 KJ/mol increase | Link to glyoxylate metabolism and ROS control^18^. | Erythrocytes |
| arginine[m] + H2O[m] => ornithine[m] + urea[m] | Arginine and proline metabolism | More than 3 KJ/mol decrease | Balance of NO and ammonia levels^37^. | Erythrocytes |
| acetyl-CoA[c] + putrescine[c] => CoA[c] + H+[c] + N-acetylputrescine[c] | Arginine and proline metabolism | More than 3 KJ/mol increase | Regulation of polyamine levels^21^. | Erythrocytes |
| 3-phosphonooxypyruvate[c] + glutamate[c] <=> 3-phosphoserine[c] + AKG[c]  3-phosphoserine[c] + H2O[c] => Pi[c] + serine[c] | Glycine, serine and threonine metabolism | Almost 3 KJ/mol increase | Precursor to serine biosynthesis^23^. | Erythrocytes |
| 3-phosphoserine[c] + H2O[c] => Pi[c] + serine[c] | Glycine, serine and threonine metabolism | More than 3 KJ/mol increase | The precursor to serine metabolism and cellular proliferation mechanism^45^. | Erythrocytes |
| serine[c] + THF[c] <=> 5,10-methylene-THF[c] + glycine[c] + H2O[c] | Glycine, serine and threonine metabolism | More than 4 KJ/mol decrease | Central to one-carbon metabolism is important for DNA synthesis and several amino acid metabolisms^23,24^. | Erythrocytes |
| serine[c] => H+[c] + NH3[c] + pyruvate[c] | Glycine, serine and threonine metabolism | More than 3 KJ/mol decrease | Balance of proper ammonia level^s37^. | Erythrocytes |
| pyruvate[c] + serine[c] => alanine[c] + hydroxypyruvate[c] | Glycine, serine and threonine metabolism | More than 3 KJ/mol increase in + ∆G^o^ | Connection to several amino acid metabolism and energy metabolism^16^ | Erythrocytes |
| serine[c] => dehydroalanine[c] + H2O[c] | Glycine, serine and threonine metabolism | More than 3 KJ/mol decrease | Important for modification of proteins and precursors to other amino acids as needed^24^. | Erythrocytes |
| pyruvate[x] + serine[x] => alanine[x] + hydroxypyruvate[x] | Glycine, serine and threonine metabolism | More than 3 KJ/mol increase in + ∆G^o^ | The interrelationship between amino acid and carbohydrate metabolism^25^. | Erythrocytes |

**References for Supplementary File 2**

1.	Ganeshan, K. & Chawla, A. Metabolic Regulation of Immune Responses. *Annu. Rev. Immunol.* **32**, 609–634 (2014).

2.	Fox, C. J., Hammerman, P. S. & Thompson, C. B. Fuel feeds function: energy metabolism and the T-cell response. *Nat. Rev. Immunol.* **5**, 844–852 (2005).

3.	DeBerardinis, R. J., Lum, J. J., Hatzivassiliou, G. & Thompson, C. B. The Biology of Cancer: Metabolic Reprogramming Fuels Cell Growth and Proliferation. *Cell Metab.* **7**, 11–20 (2008).

4.	ATP-Citrate Lyase Links Cellular Metabolism to Histone Acetylation | Science. https://www-science-org.libproxy.unl.edu/doi/full/10.1126/science.1164097.

5.	Diez, V., Traikov, S., Schmeisser, K., Adhikari, A. K. D. & Kurzchalia, T. V. Glycolate combats massive oxidative stress by restoring redox potential in Caenorhabditis elegans. *Commun. Biol.* **4**, 151 (2021).

6.	Saini, M., Nagpaul, J. P. & Amma, M. K. P. Effect of propane-1,2-diol ingestion on carbohydrate metabolism in female rat erythrocytes. *J. Appl. Toxicol.* **13**, 69–75 (1993).

7.	Miller, K. D. *et al.* Acetate acts as a metabolic immunomodulator by bolstering T-cell effector function and potentiating antitumor immunity in breast cancer. *Nat. Cancer* **4**, 1491–1507 (2023).

8.	Shiba, S. *et al.* Acetaldehyde exposure underlies functional defects in monocytes induced by excessive alcohol consumption. *Sci. Rep.* **11**, 13690 (2021).

9.	Fico, A. *et al.* 2-deoxy-d-ribose induces apoptosis by inhibiting the synthesis and increasing the efflux of glutathione. *Free Radic. Biol. Med.* **45**, 211–217 (2008).

10.	Martí i Líndez, A.-A. & Reith, W. Arginine-dependent immune responses. *Cell. Mol. Life Sci. CMLS* **78**, 5303–5324 (2021).

11.	Kim, J.-A., Kong, C.-S., Pyun, S. Y. & Kim, S.-K. Phosphorylated glucosamine inhibits the inflammatory response in LPS-stimulated PMA-differentiated THP-1 cells. *Carbohydr. Res.* **345**, 1851–1855 (2010).

12.	Hansen, A. M. & Caspi, R. R. Glutamate joins the ranks of immunomodulators. *Nat. Med.* **16**, 856 (2010).

13.	Frontiers | The role of glutamate receptors in the regulation of the tumor microenvironment. https://www.frontiersin.org/journals/immunology/articles/10.3389/fimmu.2023.1123841/full.

14.	Bogdan, C. Nitric oxide and the immune response. *Nat. Immunol.* **2**, 907–916 (2001).

15.	Wijnands, K. A., Castermans, T. M., Hommen, M. P., Meesters, D. M. & Poeze, M. Arginine and Citrulline and the Immune Response in Sepsis. *Nutrients* **7**, 1426 (2015).

16.	Arginine, ornithine and citrulline supplementation in rainbow trout: Free amino acid dynamics and gene expression responses to bacterial infection - ScienceDirect. https://www.sciencedirect.com/science/article/pii/S1050464820300267?via%3Dihub.

17.	Abusalamah, H., Reel, J. M. & Lupfer, C. R. Pyruvate affects inflammatory responses of macrophages during influenza A virus infection. *Virus Res.* **286**, 198088 (2020).

18.	Ginguay, A., Cynober, L., Curis, E. & Nicolis, I. Ornithine Aminotransferase, an Important Glutamate-Metabolizing Enzyme at the Crossroads of Multiple Metabolic Pathways. *Biology* **6**, 18 (2017).

19.	Bicarbonate enhances the inflammatory response by activating JAK/STAT signalling in LPS + IFN-γ-stimulated macrophages | The Journal of Biochemistry | Oxford Academic. https://academic-oup-com.libproxy.unl.edu/jb/article/167/6/623/5711293?login=true.

20.	Ariav, Y., Ch’ng, J. H., Christofk, H. R., Ron-Harel, N. & Erez, A. Targeting nucleotide metabolism as the nexus of viral infections, cancer, and the immune response. *Sci. Adv.* **7**, eabg6165 (2021).

21.	Frontiers | The role of polyamine metabolism in remodeling immune responses and blocking therapy within the tumor immune microenvironment. https://www.frontiersin.org/journals/immunology/articles/10.3389/fimmu.2022.912279/full.

22.	Wittmann, C. *et al.* Hydrogen Peroxide in Inflammation: Messenger, Guide, and Assassin. *Adv. Hematol.* **2012**, 541471 (2012).

23.	Serine metabolism antagonizes antiviral innate immunity by preventing ATP6V0d2-mediated YAP lysosomal degradation - ScienceDirect. https://www.sciencedirect.com/science/article/pii/S1550413121001133.

24.	Wu, Q., Chen, X., Li, J. & Sun, S. Serine and Metabolism Regulation: A Novel Mechanism in Antitumor Immunity and Senescence. *Aging Dis.* **11**, 1640 (2020).

25.	Brosnan, J. T. Glutamate, at the Interface between Amino Acid and Carbohydrate Metabolism. *J. Nutr.* **130**, 988S-990S (2000).
